# High-Molecular-Weight Genomic DNA Extraction from Recalcitrant Australian Plants: An Optimised CTAB Protocol for *Anigozanthos*

**DOI:** 10.64898/2026.08.29.741951

**Authors:** Rashika Rajput, Linkon Saha, Zeeshan Ahmed, Pia Naiker, Lien Do, Andrew Bisset, Cornelia M Hooper

## Abstract

High-phenolic plant genera present a major technical limitation in genomic research. Standard extraction approaches that perform reliably across diverse flora often perform poorly when applied to recalcitrant taxa, producing low DNA yield and integrity incompatible with sequencing requirements. The genus *Anigozanthos* (Kangaroo paws) from the family *Haemodoraceae* exemplifies this problem. We identified key physicochemical factors governing extraction failure in this genus and resolved them through targeted modifications to lysis chemistry and contaminant management. The resulting protocol achieved a near threefold improvement in DNA purity, substantially reducing contaminant carry over and consistently yielded high-integrity, long DNA fragments (DIN > 7) across a diverse sample set spanning cultivated and wild material across four diverse genera of *Haemodoraceae*. We also tested a straightforward purity assessment framework that can be implemented in any standard molecular laboratory, enabling rapid pre-submission quality assessment without the need for specialised equipment. Together these advances open a practical path to genomic characterisation of *Anigozanthos* that establishes a transferable model for genomic research across Australia’s chemically complex native flora.

## Introduction

CTAB-based genomic DNA extraction, first described by Murray and Thompson (1980) **(1)** and refined for plant molecular biology by Doyle and Doyle **(1987) (2)**, remains the primary method for isolating high-molecular-weight (HMW) DNA from Australian native plant species, including *Acacia, Telopea, Hibbertia*, and numerous orchid taxa **(3)**, as it yields DNA fragments suitable for short and mostly also long read sequencing. The Australian family members of *Haemodoraceae*, have until recently, received limited genomic attention and no publicly available long read based genomic reference exists. In our laboratory, this original protocol with some known modifications yielded high-quality genomic DNA for a number of West Australian plant species from different families in monocot and dicot orders **(4)**. However, initial extraction attempts from fresh *Anigozanthos* leaf tissue revealed extremely low yields and degraded DNA despite spectral readings promising desirable concentrations and QC assessment spectral ratios. This result is consistent with the exceptionally complex and distinct metabolic profile of the *Haemodoraceae* family sitting within recalcitrant Australian native species. Abundant secondary metabolites including polyphenols, tannins, terpenoids, and complex polysaccharides are known players in ecological responses to strong UV radiation, nutrient poor soils and herbivory pressure **(5)**.

Members of the *Haemodoraceae* family accumulate phenylphenone pigments (*e*.*g*. haemocorin, lachnanthocarpone and dihydroanigozanthin) in leaf and inflorescence tissue that are known to co-extractant during nucleic acid isolation **(6)**. Upon tissue disruption, oxidised phenolics and polysaccharides permanently bind and co-isolate with DNA, causing browning, inflate spectrophotometric estimates and reduce stability during storage, shipment and downstream enzymatic applications **(3,5,7)**. Structural and mechanical properties of sclerophyllous tissues further complicate extraction. Fibrous structure, thick cuticles, and lignified cell walls require thorough homogenisation, increasing the risk of DNA shearing and reduced integrity (DIN) **(3,8)**.

Several optimisation strategies have been described for recalcitrant plant species, including β-mercaptoethanol (2-ME) as a reducing agent. These include adding sorbitol prewashes to reduce polysaccharide content, PVP to sequester phenolics and conservative aqueous-phase recovery during chloroform separation **(3,8)**. Although effective in many taxa, their performance remains highly species dependent **(3)**. While short-read sequencing can tolerate fragmented and moderately impure DNA, long-read sequencing platforms and chromosome-level genome assembly require high-molecular-weight DNA of substantially higher purity. As these technologies become increasingly central to structural and epigenomics analyses that are required for systems level plant understanding, robust extraction protocols for chemically complex taxa such as *Anigozanthos* with increasing horticultural and ecological significance have become essential **(9)**. Here we present our optimised DNA extraction protocol that consistently yields high-quality DNA (DIN 7-9) for all tested *Anigozanthos* species and closely related genera.

### DNA extraction and Protocol Optimisation

The Paw-CTAB DNA extraction protocol was developed through three sequential optimisation phases using fresh leaf tissue from *Anigozanthos, Conostylis, Blancoa* and *Macropidia*. Tissue was selected from the basal, actively growing region of the leaf immediately above the meristem and included some of the meristem tissue.

#### Phase 1 (Initial Protocol)

Leaf tissue (50-200mg) was snap frozen in liquid nitrogen **(3)** and mechanically crushed to a fine powder using a Retsch MM 301 Mixer Mill (Retsch GmbH & Co. KG, Haan, Germany) with 50 ml stainless steel (grade 1.4112) grinding jars and one 30 mm steel grinding ball. In a subset of extractions, a sorbitol pre-wash was applied prior to reduce polysaccharide and polyphenol co-extraction **(10)**. Crushed tissue powder was washed in ice-cold sorbitol buffer, pelleted by centrifugation and the supernatant was discarded before lysis. Tissue was lysed in CTAB lysis buffer (2% CTAB, 1.4M NaCl, 20mM EDTA, 100mM Tris-HCl @ pH8.0) supplemented with 1% PVP-40k **(5)** and 0.2% 2-ME and 0.1 mg/ml RNase A (Thermo Scientific RNase A, DNase and Protease-free 10 mg, 10 mg/ml LOT-3069994, Ref-EN 0531) on the day of extraction **(1,2)**. The frozen, weighed tissue powder was added to pre-heated 400 µL of CTAB lysis buffer and incubated at 65°C for 30 mins with gentle mixing (Thermomixer, Eppendorf AG 22331, Hamburg, No. 5335). Chloroform extraction was performed adding equal volume, followed by sodium acetate and ice-cold isopropanol precipitation of the DNA in the isolated upper aqueous phase. The pellet was washed twice with 70% ethanol **(11)** and resuspended in 30 µL sterile MilliQ water or Elution Buffer (EB, Qiagen DNeasy kit).

#### Phase 2 (RNA-Contamination Reduction)

In phase 2, RNase A (Thermo Scientific RNase A, DNase and Protease-free 10 mg, 10 mg/ml LOT-3069994, Ref-EN 0531) was removed from the lysis step and instead added directly to the chloroform after lysis (10 µL of 10mg/ml; 37 °C for 15 mins) allowing enzymatic RNA removal after the bulk of co-contaminants had been separated. Tissue input was standardised to 100-150 mg, and lysis solution was increased to 700 µL (tissue to lysis solution ≤ 20 mg/100 µL), similar to Neville et al. **(2020) (4)**. Incubation temperature was reduced to 60 °C and duration of mixing increased to 60 mins to improve DNA release while minimising thermal degradation. Following lysis, an equal volume of chloroform (700 µL) was added and mixed. After centrifugation at maximum speed (4 °C, 10 mins), 500 µL of the aqueous upper phase was transferred to a new collection tube. Precipitation and washing steps remained unchanged from phase 1 with final resuspension volume of 30-50 µL.

#### Phase 3 (Chemical-Co-Contamination Reduction)

Trials of sorbitol washes in phase 1 in *Anigozanthos* samples lead to diminishing DNA yield and did not pose a solution to removing large amounts of long-chained contaminants. Instead, we introduced a clarifying centrifugation step (20,000 x g), 10 mins, 4 °C) after sufficient cell lysis and before RNase A (Monarch RNase A T3018L-20 mg/ml, NEB # T3010) treatment. This pelleted cell debris, large polysaccharide aggregates and existing RNase inhibitors that showed to interfere with RNA digestions in previous protocols. The clarified supernatant was then collected for chloroform 2-phase extraction. In addition, an increase of 2-ME concentration from 0.2% to 0.5% was required which is known to reduce oxidation of thiol groups and disulphide bonds **(3,8)**, protecting *Anigozanthos* DNA from high phenolic-driven damage during lysis. A second chloroform extraction was incorporated to improve purity, and pellet recovery centrifugation time was extended to 40 mins. The pelleted DNA was resuspended in 50 µL sterile MilliQ H_2_O or EB (Elution Buffer, Qiagen DNeasy kit) and stored at −20 °C. Comprehensive detail for extraction buffer compositions, reagent volumes and workflow are provided in the final protocol **(25)**.

#### Quality control assessment

Extracted DNA samples were assessed using a Nanodrop One spectrophotometer (Thermo Scientific) and a Qubit 3 Fluorometer (Invitrogen, Thermo Fisher Scientific) with the Qubit dsDNA Broad Range Assay Kit (Q32853, Thermo Fisher Scientific). The ratio of DNA concentration derived from Nanodrop to Qubit (Nanodrop–Qubit ratio (N/Q)) was calculated for each sample and used as a practical indicator of phenolic co-contamination, alongside A260/280 and A260/230 ratios provided directly by the Nanodrop spectrophotometer. DNA yield (ng/mg fresh tissue) was calculated as Qubit concentration (ng/µL) × elution volume (µL) ÷ tissue weight (mg). Not all samples yielded sufficient material for every quality assessment; as a result, sample numbers vary across figures: Batch 1 is represented by n=35 for DIN-based analyses (Figure 2) and n=26 for N/Q analyses (Figures 1 and 3), reflecting the number of samples for which both Nanodrop and Qubit measurements were available. Samples submitted for Illumina short-read sequencing (150 bp paired-end) were assessed by gDNA TapeStation analysis (Agilent 4200 TapeStation System, Agilent Technologies) performed by Macrogen.

**Figure 1.**
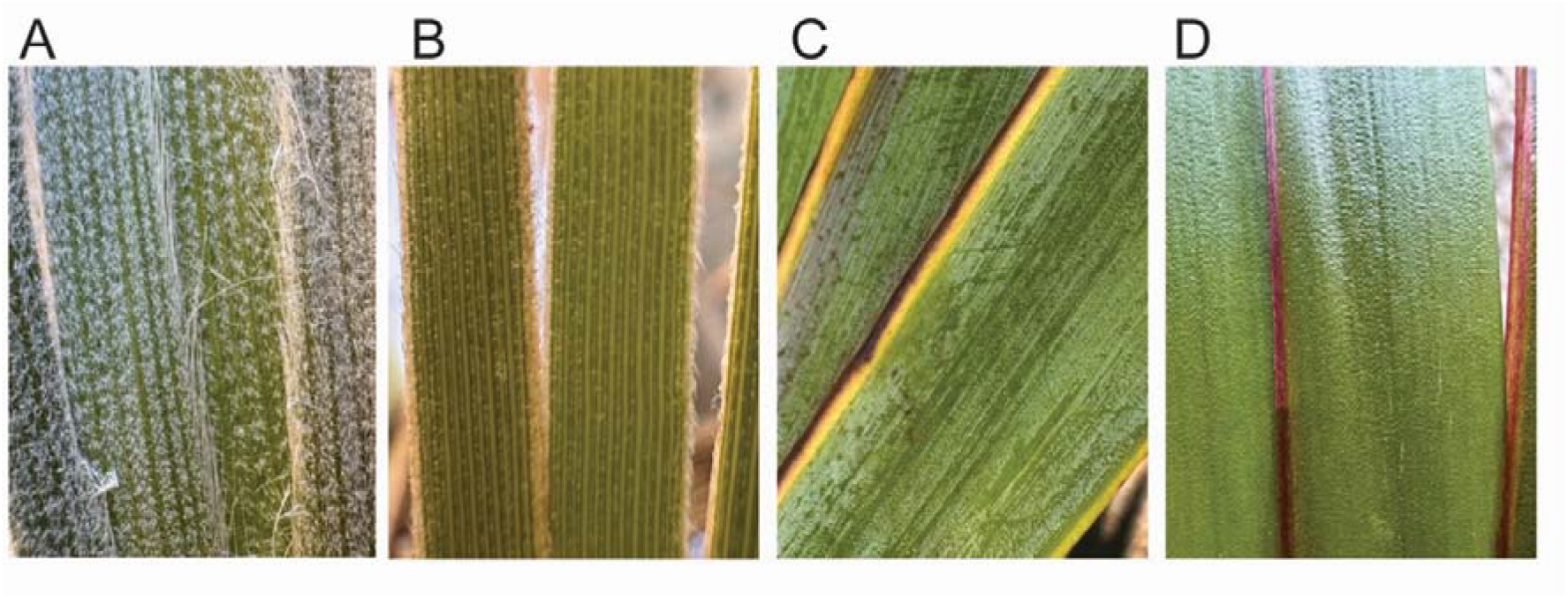
Representative macro-photographs of leaf surface texture in *Anigozanthos* and three allied *Haemodoraceae* genera examined in this study, illustrating the morphological diversity relevant to extraction difficulty across the family. Plant material was sourced from Zanthorrea Nursery, Western Australia. **(A)** *Conostylis candicans*, showing the dense, hairy leaf surface characteristic of the genus. **(B)** *Blancoa canescens*, showing the strappy, grey-green foliage habit. **(C)** *Macropidia fuliginosa*, showing the waxy, glaucous leaf surface. **(D)** *Anigozanthos flavidus* (early spring growth), showing the fleshy, waxy, and glossy leaf surface typical of the species. Photographs: Cornelia M. Hooper

**Figure 2.**
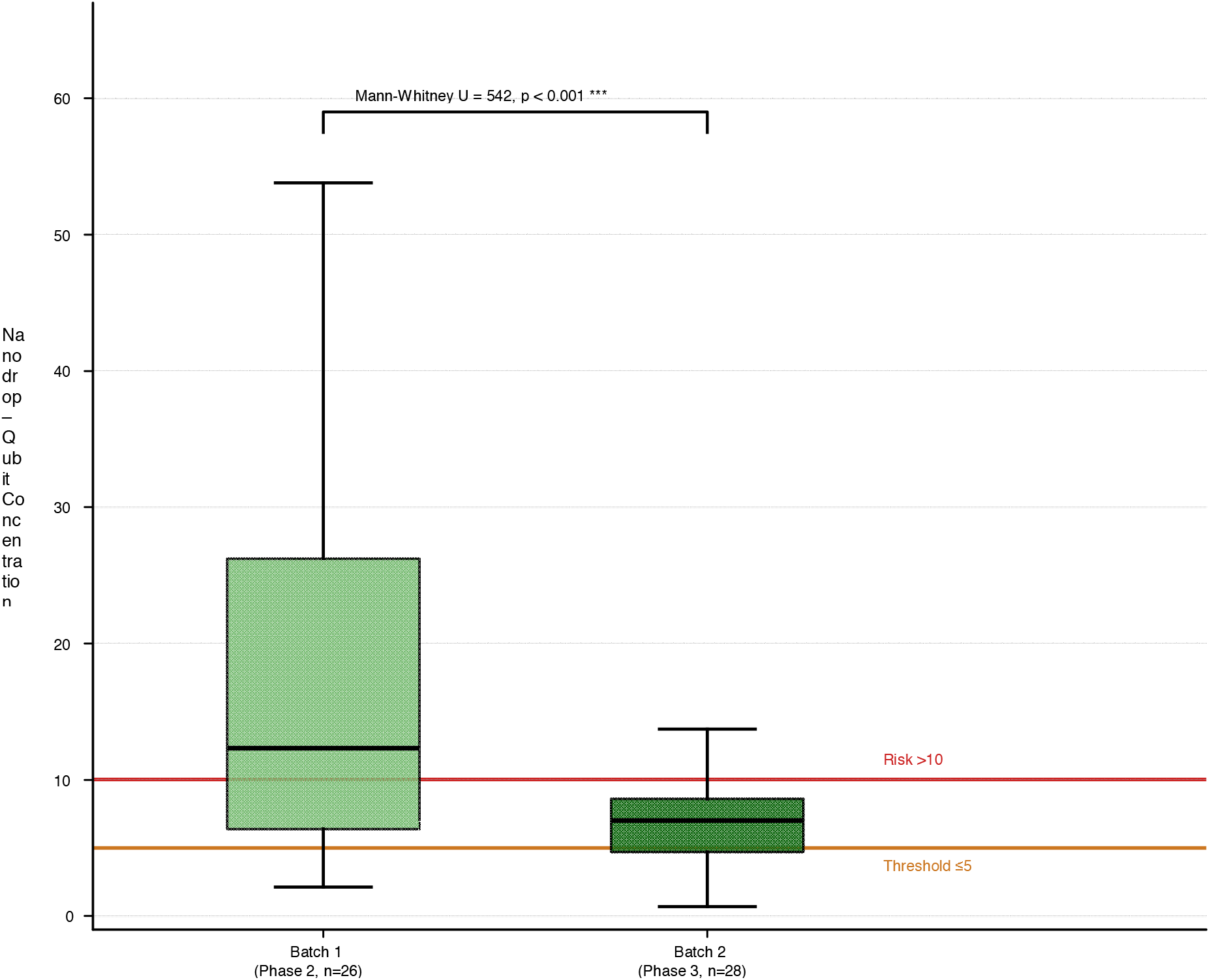
Nanodrop–Qubit concentration ratio (N/Q) for Batch 1 (Phase 2, n=26, light green) and Batch 2 (Phase 3, n=28, dark green). Dashed orange line: acceptance threshold ≤5. Dashed red line: contamination risk threshold >10. Mann-Whitney U = 542, p < 0.001. The 2.8-fold reduction demonstrates substantially reduced phenolic co-extraction using the Paw-CTAB protocol.

**Figure 3.**
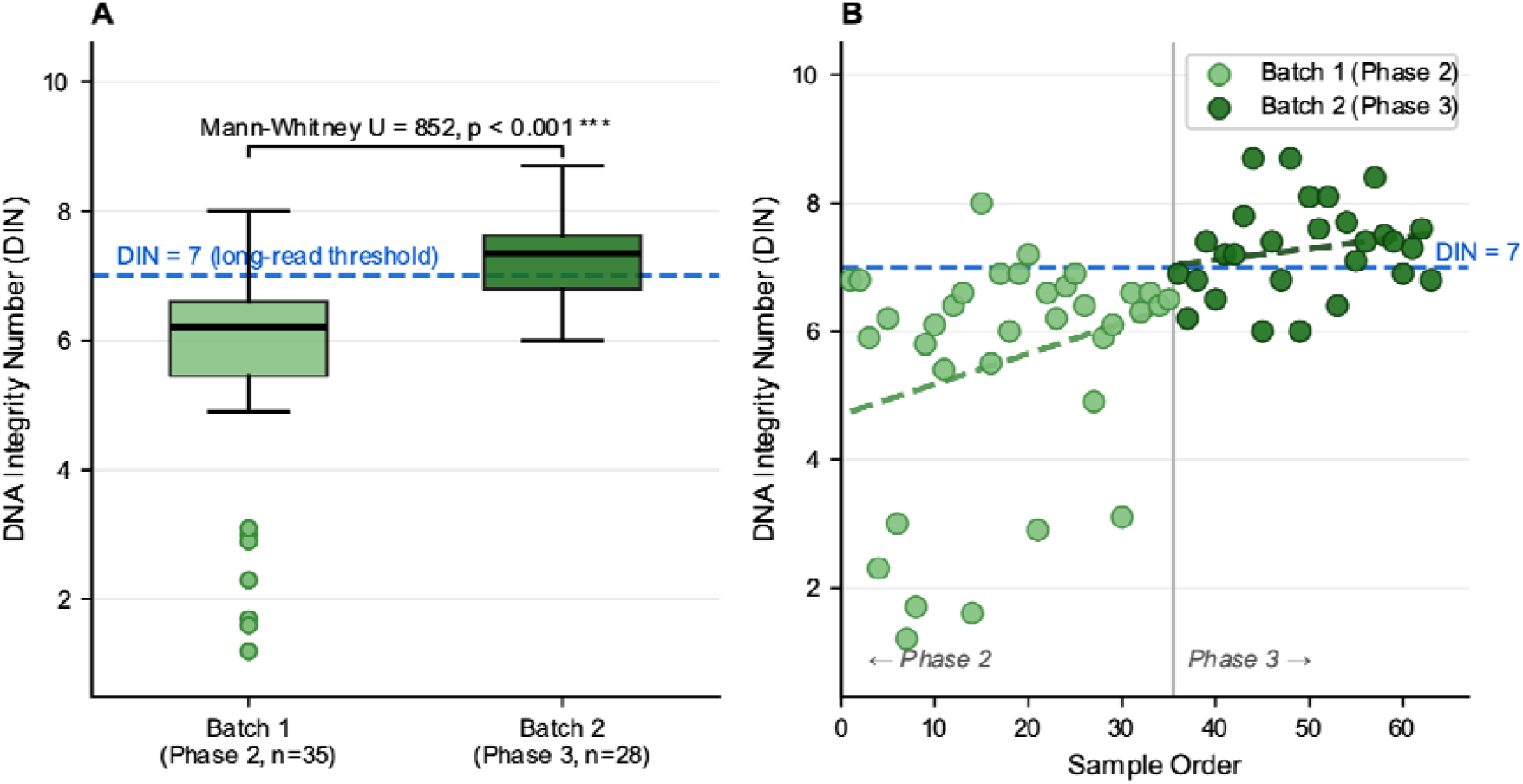
DNA Integrity Number (DIN) for Batch 1 (Phase 2, n=35, light green) and Batch 2 (Phase 3, n=28, dark green). **(A)** Box plot comparison; Batch 2 shows a significantly higher median and substantially reduced variance. Mann-Whitney U = 852, p < 0.001. Blue dashed line: DIN = 7 threshold for long-read library construction. **(B)** Sample-wise DIN values across extraction order with linear trend lines (dashed), showing the consistent increase of the DNA integrity with protocol optimisation within and across batches.

#### Data analysis

Statistical comparison between Batch 1 and Batch 2 was performed using Mann-Whitney U test, as DIN and Nanodrop-Qubit ratio data were non-normally distributed. Tests were one - tailed for DIN and ratio and twotailed for yield.

Sequencing was performed on samples submitted across two separate batches. Batch 1 contained samples prepared using versions of phase 1 and phase 2 protocols in iterative improvement steps. The optimised phase 3 method “Paw-CTAB” was applied to all Batch 2 sample set comprising 28 accessions spanning *Anigozanthos* cultivars and wild accessions, and the allied genera *Macropidia* (n= 2), *Conostylis* (n=1) and *Blancoa* (n=2).

## Results

### Poor DNA recovery and stability in *Haemodoraceae* despite modified protocols for recalcitrant plants

The CTAB method of Doyle and Doyle **(1987) (2)**, as adapted by Neville et al. **(2020) (4)**, has demonstrated reliable performance across a range of Australian plant taxa. Across mixed Australian native taxa, this method alone outperformed modified protocols with additional sorbitol pre-washes as well as a number of commercial kits delivering DNA yields ranging 230-880 ng/mg fresh weight **(12,13)**. All protocols were able to recover sufficient DNA for Illumina sequencing for the mix of Australian natives not including *Haemodoraceae*. Notably, the introduction of a sorbitol pre-wash intended to reduce polyphenol load paradoxically produced inconsistent A260/230 ratios and substantially reduced yield. Across 19 Australian frozen seagrass samples from multiple Western Australian collection sites, CTAB similarly outperformed the Qiagen DNeasy Plant Pro Mini Kit, achieving yields of 2.6– 119.5 ng/mg. The wide range in CTAB yield reflected inter-sample biological variation across species and collection sites rather than protocol inconsistency with N/Q of 1.8–2.01, compared with 5.6–31.3 ng/mg and ratios of 0.05–1.19 for the commercial kit **(14)**.

*Anigozanthos* and allied *Haemodoraceae* genera present distinct tissue properties that complicate DNA extraction (Figure 1). The leaves are fibrous and tough with thick, waxy and glossy surfaces (Figure 1C, 1D) that resist mechanical homogenisation and repel aqueous lysis buffer. Upon disruption, the homogenate browns rapidly, indicating immediate phenolic oxidation consistent with the high phenylphenalenone content characteristic of this family **(6)**. These properties are shared across the allied genera examined in Figure 1A, 1B, 1C suggesting a family- wide extraction challenge rather than a speciesspecific one.

Considering the above performance in numerous Australian taxa, the original CTAB **(4)**, (phase 1) produced surprisingly low yields using fresh *Anigozanthos* tissue. In fact, a wide range of commercial kits trialled in comparison to the original CTAB protocol all yielded less than 2 ng/mg by Qubit fluorometric quantification, accompanied by Nanodrop readings one to two orders of magnitude higher. This mismatch of concentration reading between Nanodrop and Qubit methods is often indicative of severe phenolic co-contamination **(15)**. The surprising failure of all approaches poses the question of a *Haemodoraceae* or species-specific extraction challenge. However, out of all trialled methods the CTAB approach seemed to be the best starting point for systematic optimisation of the chemistry. We extracted DNA from freshly cut leaf tissue using incremental steps of the phase 2 protocol (Batch 1, n=35) and then the fully optimised Paw-CTAB (Batch 2, n=28). While not all samples were sent for TapeStation assessment and sequencing, all were assessed by Qubit and Nanodrop.

Batch 1 (phase 2 protocol) yields measured by Qubit averaged 14.5 ± 10.1 ng/mg fresh tissue which is low but seemed stable for this genus. Nanodrop-derived yields were markedly higher at 164.5 ± 95.8 ng/mg, producing a mean N/Q of 18.5 ± 16.0 (range 2.1– 53.8) likely due to impurities such as phenolic co-extractants that increase absorbance reading in the Nanodrop measurement but not in the probe-specific Qubit **(Figure 2)**. A260/280 Nanodrop values were within the normal range for plant DNA (mean 1.98 ± 0.11), as is typical even in phenolic-contaminated samples; however, A260/230 ratios were highly variable (mean 1.72 ± 0.33, range 1.0–2.3), with many samples below the 1.8 acceptability threshold. TapeStation analysis revealed a mean DIN of 5.55 ± 1.79 ranging from 1.2–8.0 **(Figure 3A)**, with 33/35 samples not meeting the DIN ≥7 threshold required for long-read library construction with 13 failing, 20 placed on hold, and only 2 passing. Critically, several samples showed reduced DNA concentration and fragment length upon arrival at the sequencing facility compared with in-house measurements indicating ongoing DNA damage as is characteristic for chemical contaminants **(16)**.

Batch 2 (Paw-CTAB) yields improved to a mean of 20.6 ± 11.7 ng/mg. The mean N/Q fell to 6.5 ± 3.4 (range 0.7–13.7), a 2.8-fold reduction overall as well as more consistency relative to Batch 1 **(Figure 2)**. Minimal degradation between in-house assessment and the sequencing provider was observed in any Batch 2 sample, confirming that the removal of suspected phenolic co-contaminants stabilised the extracted DNA. TapeStation analysis confirmed a mean DIN of 7.28 ± 0.73 (range 6.0–8.7) **(Figure 3A)**, with all 28 samples successfully generating sequencing libraries. Out of all samples, 18 samples (64%) met the DIN ≥7 threshold for long - read library construction. The remaining 10 samples have DIN values of 6.0 – 6.9, narrowly below the threshold, with no association found with tissue weight, N/Q, or extraction date, suggesting normal biological variation between accessions rather than an unresolved protocol limitation.

### phase 2 is sufficient for short reads while the phase 3 paw-CTAB is more suited to long read sequencing

The minimum DIN requirement for long-read sequencing library construction (TruSeq Nano DNA, PacBio SMRT, Oxford Nanopore) is DIN ≥7, with a minimum total DNA input of ≥1 μg **(17)**. Under the phase 2 partially modified CTAB protocol, only 2 of 35 samples met this threshold. The remaining 33 samples (94%) were below, unsuitable for long-read applications. Comparing in-house Qubit concentrations (mean 80.3 ng/µL) with TapeStation concentrations measured by Macrogen on arrival (mean 79.0 ng/µL) showed no statistically significant change during transit (Wilcoxon signed-rank test, p = 0.65, n=39). We therefore conclude that the quality control failures observed in Batch 1 were primarily driven by residual co-extracted contaminants rather than DNA loss during shipment **(5, 16)**. Although these contaminants are unlikely to substantially reduce the total amount of measurable double-stranded DNA, they can promote progressive fragmentation of HMW DNA during storage and transport. Consequently, Qubit-derived DNA concentrations remain largely unchanged while DNA integrity (DIN) declines, reducing suitability for long-read sequencing. In contrast, the Paw-CTAB, 18 of 28 samples (64%) achieved DIN ≥7 while the remaining samples had DIN values only marginally below this threshold (6.0 – 6.9) **(Figure 3B)**. No significant change in DNA concentration was observed following shipment, indicating that sample integrity was maintained.

Samples with DIN >3 from both batches successfully generated 300 bp Illumina sequencing libraries. In contrast, the Paw-CTAB protocol consistently produced DNA that met, or was only marginally below, the accepted DIN threshold for long-read library preparation. While DNA concentration remained stable following shipment, DNA integrity was more sensitive to the effects of residual co-extracted contaminants, supporting the importance of minimising contamination when preparing HMW DNA for long-read sequencing.

### N/Q provide a simple cost-effective pre-submission quality indicator

Because co-extracted polysaccharides and phenolics inflate UV absorbance at 260 nm, Nanodrop measurements overestimate DNA concentration in contaminated samples. In contrast, the Qubit fluorescent dye binds specifically to double-stranded DNA (dsDNA) **(15)**, providing a more accurate estimate of DNA concentration. For pure DNA, the Nanodrop and Qubit measurements are expected to agree, producing a N/Q close to 1. Increasing divergence between the two measurements therefore provides a practical indicator of coextracted contaminants. This simple ratio has proven to be a rapid and cost-effective quality control measure that complements DNA integrity assessment by agarose gel electrophoresis or TapeStation analysis. Whereas DIN reflects the integrity of DNA at the time of measurement, the N/Q provides an indication of contaminant carry-over that is not captured by fragment analysis alone and may influence DNA stability during storage and transport.

Using the Paw-CTAB reduced the mean ratio from 18.5 (Batch 1) to 6.5 (Batch 2) (Mann-Whitney U = 542, p < 0.001; **Figure 2**). Batch 2 samples with ratios below 5 passed DIN quality control without exception **(Figure 2)**. Batch 1 samples with ratios exceeding 10 consistently failed or showed degradation **(Figure 2)**. Masago et al. **(15)** confirmed that Nanodrop systematically overestimates DNA concentration relative to Qubit fluorometry across tissue types, consistent with our observations. Porebski et al. **(5)** established that high phenolics and polysaccharide content in plant tissue interferes with spectrophotometric DNA quantification. Building on these findings and our experimental data, we propose a ratio of ≤5 as a practical pre-submission acceptance criterion for highly phenolic *Haemodoraceae*, with values of 1-3 providing the highest confidence of sequence success. DNA integrity (DIN) was significantly higher in Batch 2 (median 7.3) than Batch 1 (median 6.2; U = 852, p < 0.001; **Figure 3A**). The N/Q was significantly lower in Batch 2 (median 6.9) than Batch 1 (median 13.0; U = 542, p < 0.001; **Figure 2)**, indicating reduced phenolic contamination. DNA yield was also significantly higher in Batch 2 (median 19.8 ng/mg) than Batch 1 (median 13.5 ng/mg; U = 697, p = 0.012). Across all samples, the N/Q was negatively correlated with DNA integrity (Spearman ρ = −0.35, p = 0.009; **(Figure 4)**. Samples with lower N/Q generally exhibited higher DIN values, consistent with reduced contaminant carry-over following protocol optimisation. The N/Q reflects residual phenolic contamination, whereas DIN reflects its effect on DNA integrity. Together, these metrics provide a more complete assessment of DNA quality **(18)**. Together these results confirm that the Paw-CTAB produced robust improvements across all dimensions of DNA quality relative to the phase 2 protocol.

**Figure 4.**
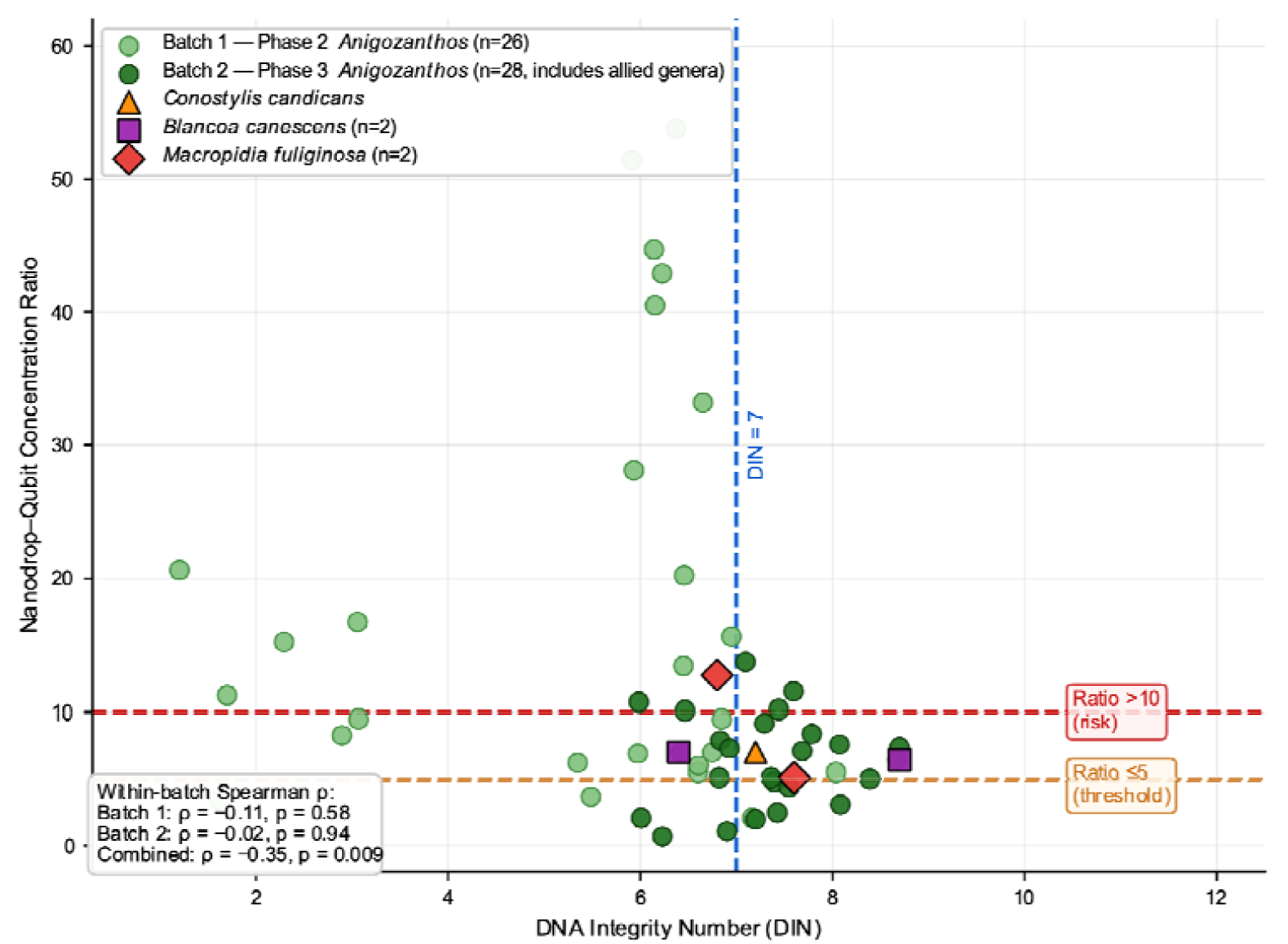
Nanodrop–Qubit ratio (N/Q) versus DNA integrity Number (DIN) for Batch 1 (Phase 2, n=26, light green circles) and Batch 2 (Phase 3, n=28, dark green circles). Batch 2 includes 3 other *Haemodoraceae* genera indicated by distinct symbols: *Conostylis candicans* (orange triangle), *Blancoa canescens* (purple squares, n=2), and *Macropidia fuliginosa* (red diamonds, n=2). Dashed orange: ratio threshold ≤5. Dashed red: risk threshold >10. Blue dashed: DIN = 7. Spearman correlation statistics (inset, bottom left) show within-batch and combined correlations demonstrating that both metrics are needed for a complete pre-submission quality assessment.

### Performance of Paw-CTAB is consistent across allied *Haemodoraceae* genera

The Paw-CTAB was applied mainly to the genus *Anigozanthos* but also to 3 specimen from closely *Haemodoraceae* genera in Batch 2, namely *Macropidia fuliginosa* (n=2, DIN=7.6 and 6.8, N/Q of 5.1 and 12.7), *Conostylis candicans* (DIN=7.2, N/Q=7.0) and *Blancoa canescens* (n=2, DIN=8.7 and 6.4, N/Q=6.4 and 7.0) (Figure 4). Successful DNA recovery was achieved in four of the six recognised *Haemodoraceae* genera, including extensive testing across *Anigozanthos*. Together, these results suggest that the Paw-CTAB protocol is broadly applicable across the *Haemodoraceae*.

## Discussion

This report highlights the considerable difficulty of extracting high integrity gDNA from *Anigozanthos* using both commercial kits and the standard CTAB method. High-quality *Anigozanthos* gDNA (DIN 7–9) was consistently achieved using our modified CTAB-based approach (phase 3) called Paw-CTAB with DNA quality governed principally by effective tissue disruption, standardised tissue input relative to CTAB lysis buffer volume (≤20 mg per 100 µL) and systematic removal of debris and contaminants prior to phase separation. Together these improvements prevented buffer overload and minimised co-precipitation of contaminants with gDNA, the two significant drawbacks observed when working with high-phenolic Australian native plant species **(3,5,8)**.

The key optimisation was balancing chemical ratios in the secondary metabolite-rich tissues. This included tissue input standardisation to not exceed the CTAB buffer’s detergent and chelating capacity, leaving oxidised phenolics free that are known to bind and co-precipitate with DNA **(3,19)**. The initial extra centrifugation removes excess debris that interferes with RNase activity and depletes the effective capacity of the extraction chemistry, an often-noted problem **(4,19)**. For *Haemodoraceae*, the sorbitol pre-wash is ineffective, and its negative effect on yield indicates phenolic co-release as the primary constraint on extraction. Residual phenolic compounds pose a high risk beyond the extraction itself, as they can drive ongoing oxidative DNA degradation during storage and transit **(3,5)**. This explains the systematic discrepancy observed in Batch 1, where several samples show acceptable in-house integrity but failed quality control at the sequencing provider **(17)**. The Nanodrop-Qubit ratio served as a practical purity indicator, as the Nanodrop A260 measurements are inflated by phenolics, whereas Qubit more selectively quantifies dsDNA via a fluorescent dye largely insensitive to these contaminants **(15)**. Therefore, the N/Q ratio provides a rapid proxy for extract purity useful as a pre-sequencing quality screen in routine molecular workflows without the need for TapeStation.

## Conclusion

Molecular studies of the *Haemodoraceae* have provided important insights into the evolution, taxonomy and conservation of the family but have been limited by the availability of high-quality genomic resources **(20,21,22)**. The Paw-CTAB protocol resolves the primary barrier to genomic DNA extraction from *Anigozanthos* and allied *Haemodoraceae*. Consistent performance across four genera (*Anigozanthos, Macropidia, Conostylis*, and *Blancoa*) suggests the protocol is broadly applicable across the *Haemodoraceae* family. The N/Q ratio provide a practical pre-submission indicator that can be implemented in any standard molecular laboratory without TapeStation instrumentation. We recommend a N/Q ratio of ≤ 5 as an acceptance criterion for long-read sequencing applications. The continued expansion of genomic resources will underpin the study, breeding and conservation of Australian native plant species **(23,24)**. By overcoming a major bottleneck in obtaining sequencing-grade DNA, the Paw-CTAB protocol and Nanodrop–Qubit quality assessment framework unlock these opportunities for the *Haemodoraceae*.

## Acknowledgement

The authors gratefully acknowledge Digby Growns and Dr. Praful Umaretiya (Kings Park & Botanic Garden; Department of Biodiversity, Conservation & Attractions, WA) and Zanthorrea Nursery for provision of plant material used in this report. We thank Dr. David Field (Macquarie University) and Prof. Ian Small (The University of Western Australia) for their valuable contribution, scientific direction and constructive feedback throughout the development of this report. This work was supported by Australian Research Council (IE230100040).

## Author Contributions

R.R. developed and optimised the extraction protocol across phase 3, performed all laboratory investigations, carried out formal analysis and data curation, and wrote the manuscript. L.S. contributed to Phase 2 protocol optimisation and laboratory investigation. Z.A. independently verified the final method on *Anigozanthos* samples. P.N., L.D. and A.B. provided baseline extraction data from unpublished master’s thesis work that informed protocol development. C.M.H. conceived and supervised the project, acquired funding, provided plant material and macro-photographs, and revised the manuscript. All authors approved the final version of the manuscript.

## Notes

### Competing Interest Statement

The authors have declared no competing interest.

