## Supplementary material for "High-Molecular-Weight Genomic DNA Extraction from Recalcitrant Australian Plants: An Optimised CTAB Protocol for *Anigozanthos*": S1

### Batch 1

| Species / Cultivar | Genus | Tissue wt (mg) | Q20 (%) | Q30 (%) | Qubit yield (ng/mg) | Nanodrop yield (ng/mg) | N/Q ratio (ND yield ÷ Qubit yield) | TapeStation conc. (ng/µL) | DIN |
|---|---|---|---|---|---|---|---|---|---|
| Stripie | *Anigozanthos* | NULL | 98.5 | 95.8 | NULL | NULL | **NULL** | NULL | NULL |
| Early Spring | *Anigozanthos* | NULL | 98.5 | 95.3 | NULL | NULL | **NULL** | NULL | NULL |
| Anigozanthos flavidus (green form) | *Anigozanthos* | 141.3 | 98.5 | 96.5 | 17.18 | NULL | **NULL** | 23.64 | 6.8 |
| Anigozanthos 'Bush Tenacity' | *Anigozanthos* | 128.9 | 98.5 | 96.6 | 15.73 | 110.8 | **7.05** | 106.61 | 6.8 |
| Anigozanthos humilis | *Anigozanthos* | 150.0 | 98.5 | 96.6 | 11.93 | 334.9 | **28.07** | 39.34 | 5.9 |
| Anigozanthos rufus 'Frosty Yellow' | *Anigozanthos* | 145.0 | 98.5 | 96.8 | 19.54 | 297.2 | **15.21** | 68.84 | 2.3 |
| Anigozanthos kalbarriensis | *Anigozanthos* | 150.0 | 98.5 | 96.7 | 10.37 | 444.4 | **42.85** | 38.95 | 6.2 |
| Anigozanthos bicolor ssp. bicolor | *Anigozanthos* | 126.0 | 98.5 | 96.8 | 7.56 | 126.3 | **16.70** | 40.43 | 3.0 |
| Anigozanthos viridis ssp. terraspectans | *Anigozanthos* | 117.6 | 98.5 | 96.7 | 10.78 | 222.6 | **20.65** | 35.69 | 1.2 |
| Anigozanthos viridis ssp. cataby | *Anigozanthos* | 123.7 | 98.5 | 96.6 | 21.58 | 241.2 | **11.18** | 68.84 | 1.7 |
| Macropidia fuliginosa | *Macropidia* | 135.2 | 98.5 | 96.5 | 8.37 | NULL | **NULL** | 48.33 | 5.8 |
| Conostylis setosa | *Conostylis* | 130.1 | 98.5 | 95.9 | 49.21 | NULL | **NULL** | 248.95 | 6.1 |
| Anigozanthos 'Bush Ballad' | *Anigozanthos* | 157.7 | 98.5 | 96.7 | 11.64 | 72.3 | **6.21** | 38.81 | 5.4 |
| Anigozanthos 'Bush Blitz' | *Anigozanthos* | 139.6 | 98.5 | 96.6 | 5.34 | 287.1 | **53.76** | 29.17 | 6.4 |
| Anigozanthos 'Bush Bonanza' | *Anigozanthos* | 142.5 | 98.5 | 96.6 | 24.36 | 135.1 | **5.54** | 102.02 | 6.6 |
| Anigozanthos 'Bush Crystal' | *Anigozanthos* | 128.5 | 98.5 | NULL | 36.75 | 125.6 | **3.42** | 85.67 | 1.6 |
| Anigozanthos 'Bush Diamond' | *Anigozanthos* | 153.3 | 98.5 | 96.7 | 18.68 | 102.2 | **5.47** | 49.98 | 8.0 |
| Anigozanthos 'Bush Fire' | *Anigozanthos* | 172.0 | 98.5 | 96.6 | 18.49 | 67.8 | **3.67** | 72.44 | 5.5 |
| Anigozanthos 'Bush Fury' | *Anigozanthos* | 153.2 | 98.5 | 96.8 | 12.79 | 121.5 | **9.50** | 39.68 | 6.9 |
| Anigozanthos 'Bush Glow' | *Anigozanthos* | 139.0 | 98.5 | 96.7 | 19.29 | 133.7 | **6.93** | 79.97 | 6.0 |
| Anigozanthos 'Bush Inferno' | *Anigozanthos* | 153.5 | 98.5 | 96.4 | 6.46 | 100.9 | **15.62** | 36.32 | 6.9 |
| Anigozanthos 'Bush Pearl' | *Anigozanthos* | 133.3 | 98.5 | 96.8 | 21.25 | 44.8 | **2.11** | 28.91 | 7.2 |
| Anigozanthos 'Bush Surprise' | *Anigozanthos* | 133.4 | 98.5 | NULL | 19.79 | 164.9 | **8.33** | 97.67 | 2.9 |
| Anigozanthos 'Bush Zest' | *Anigozanthos* | 150.5 | 98.5 | 96.6 | 5.66 | 187.9 | **33.21** | 48.53 | 6.6 |
| Anigozanthos 'Kings Park Royale' | *Anigozanthos* | 135.7 | 98.5 | 96.7 | 3.80 | 169.9 | **44.72** | 17.85 | 6.2 |
| Anigozanthos 'Aussie Spirit' | *Anigozanthos* | 140.5 | 98.5 | 96.5 | 3.89 | NULL | **NULL** | 23.19 | 6.7 |
| Anigozanthos 'Masquerade' | *Anigozanthos* | 157.2 | 98.5 | 96.7 | NULL | 46.5 | **NULL** | 40.43 | 6.9 |
| Anigozanthos 'Big Red' | *Anigozanthos* | 156.9 | 98.5 | 96.6 | 7.03 | NULL | **NULL** | 52.04 | 6.4 |
| Anigozanthos 'Yellow Gem' | *Anigozanthos* | 145.0 | 98.5 | 96.5 | 17.26 | NULL | **NULL** | 74.00 | 4.9 |
| Anigozanthos 'Landscape Orange' | *Anigozanthos* | 157.4 | 98.5 | 96.5 | 4.38 | 225.3 | **51.43** | 29.59 | 5.9 |
| Anigozanthos 'Nugget' | *Anigozanthos* | 155.7 | 98.5 | 96.3 | 4.23 | 171.2 | **40.48** | 18.12 | 6.1 |

| | | | | | | | | | |
|---|---|---|---|---|---|---|---|---|---|
| Anigozanthos A20/100H F1 (hybrid) | *Anigozanthos* | 150.0 | 98.5 | 96.5 | 9.48 | 90.4 | **9.53** | 34.26 | 3.1 |
| Anigozanthos A20/200 F2 (hybrid) | *Anigozanthos* | 150.0 | 98.5 | 96.5 | 10.27 | 61.3 | **5.97** | 66.52 | 6.6 |
| Anigozanthos A200 (hybrid) | *Anigozanthos* | 200.0 | 98.5 | 96.6 | 32.52 | NULL | **NULL** | 360.61 | 6.3 |
| Anigozanthos E212 (hybrid) | *Anigozanthos* | 212.0 | 98.5 | 96.6 | 15.36 | NULL | **NULL** | 133.37 | 6.6 |
| Anigozanthos 'Z23/015' | *Anigozanthos* | NULL | 98.5 | NULL | NULL | NULL | **NULL** | 47.96 | NULL |
| Anigozanthos 'Z22/96' | *Anigozanthos* | NULL | 98.5 | NULL | NULL | NULL | **NULL** | 57.85 | NULL |
| Anigozanthos 'Landscape Gold' | *Anigozanthos* | 152.2 | 98.5 | 96.8 | 4.88 | 65.5 | **13.43** | 41.43 | 6.4 |
| Anigozanthos 'Bush Elegance' | *Anigozanthos* | 100.0 | 98.5 | 96.7 | 8.59 | 173.3 | **20.18** | 19.79 | 6.5 |
| Anigozanthos 'Z23/015' | *Anigozanthos* | NULL | 98.5 | 96.2 | NULL | NULL | **NULL** | 290.33 | NULL |
| Anigozanthos 'Z22/96' | *Anigozanthos* | NULL | 98.5 | 96.8 | NULL | NULL | **NULL** | 147.24 | NULL |

### Batch 2

| Species / Cultivar | Genus | Tissue wt (mg) | Q20 (%) | Q30 (%) | Qubit yield (ng/mg) | "N/Q ratio (ND yield ÷ Qubit yield) | "TapeStation conc. (ng/µL)" | DIN |
|---|---|---|---|---|---|---|---|---|
| A. flavidus Stripie | *Anigozanthos* | 144.2 | 98.7 | 94.2 | 3.50 | **1.10** | 11.90 | 6.9 |
| A. flavidus Stripie (White) | *Anigozanthos* | 170.0 | 98.8 | 94.7 | 5.60 | **0.70** | 9.71 | 6.2 |
| A. Bush Coral | *Anigozanthos* | 182.4 | 98.7 | 94.5 | 19.70 | **5.10** | 40.48 | 6.8 |
| A. Gold Velvet | *Anigozanthos* | 133.4 | 98.8 | 94.6 | 26.80 | **4.80** | 31.09 | 7.4 |
| A. manglesii | *Anigozanthos* | 163.0 | 98.8 | 94.5 | 18.80 | **10.10** | 29.40 | 6.5 |
| A. onycis | *Anigozanthos* | 157.1 | 98.7 | 94.6 | 5.30 | **2.00** | 21.29 | 7.2 |
| C. candicans | *Conostylis* | 189.0 | 98.6 | 94.3 | 23.80 | **7.00** | 194.82 | 7.2 |
| Pink Beauty | *Anigozanthos* | 138.7 | 98.8 | 94.8 | 18.10 | **8.40** | 54.93 | 7.8 |
| A. flavidus ‘Tall Orange’ | *Anigozanthos* | 159.1 | 98.7 | 94.4 | 15.90 | **7.40** | 55.11 | 8.7 |
| A. manglesii (herbarium sample) | *Anigozanthos* | 174.2 | 98.8 | 94.9 | 10.50 | **10.70** | 55.40 | 6.0 |
| A. rufus ‘Scorches Flame’ | *Anigozanthos* | 184.4 | 98.7 | 94.4 | 23.80 | **10.20** | 106.90 | 7.4 |
| A. flavidus (red) | *Anigozanthos* | 168.4 | 98.8 | 94.7 | 15.20 | **7.90** | 45.20 | 6.8 |
| B. canescens | *Blancoa* | 153.9 | 98.7 | 94.4 | 8.30 | **6.40** | 64.24 | 8.7 |
| Kings Park Royale | *Anigozanthos* | 158.9 | 98.8 | 94.5 | 22.20 | **2.10** | 27.82 | 6.0 |
| A. rufus ‘Frosty Red’ | *Anigozanthos* | 154.3 | 98.8 | 94.9 | 13.20 | **7.60** | 42.24 | 8.1 |
| A. celebration range ‘Carnivale’ | *Anigozanthos* | 156.0 | 98.7 | 94.5 | 13.30 | **11.50** | 59.23 | 7.6 |
| A. tango | *Anigozanthos* | 157.3 | 98.7 | 94.3 | 29.10 | **3.10** | 30.76 | 8.1 |
| B. canescens ‘Zan’ | *Blancoa* | 160.2 | 98.6 | 94.3 | 18.30 | **7.00** | 105.44 | 6.4 |
| A. pulcherrimus | *Anigozanthos* | 159.0 | 98.7 | 94.5 | 20.30 | **7.10** | 46.30 | 7.7 |
| A. celebration range ‘Firework’ | *Anigozanthos* | 181.4 | 98.8 | 94.9 | 20.40 | **13.70** | 62.11 | 7.1 |
| A. Bush Flare | *Anigozanthos* | 157.1 | 98.7 | 94.3 | 23.10 | **2.50** | 29.11 | 7.4 |
| A. Hybrid Orange Cross | *Anigozanthos* | 156.4 | 98.8 | 94.8 | 34.80 | **5.00** | 62.49 | 8.4 |
| A. celebration range ‘Jazz’ | *Anigozanthos* | 153.6 | 98.8 | 94.7 | 54.00 | **4.40** | 44.57 | 7.5 |
| A. Devil | *Anigozanthos* | 155.3 | 98.7 | 94.5 | 37.00 | **5.10** | 68.28 | 7.4 |
| A. landscape scarlet | *Anigozanthos* | 159.8 | 98.7 | 94.3 | 26.00 | **7.30** | 50.60 | 6.9 |
| A. Bush Pearl | *Anigozanthos* | 163.9 | 98.7 | 94.6 | 22.10 | **9.20** | 56.83 | 7.3 |
| M. fuliginosa | *Macropidia* | 154.0 | 98.7 | 94.4 | 47.70 | **5.10** | 47.89 | 7.6 |
| M. fuliginosa (dwarf) | *Macropidia* | 160.1 | 98.8 | 94.7 | 24.40 | **12.70** | 78.86 | 6.8 |
| A. Stripie White | *Anigozanthos* | 149.5 | NULL | NULL | 4.70 | **3.20** | NULL | NULL |
| A. onycis | *Anigozanthos* | 115.6 | NULL | NULL | 12.80 | **7.90** | NULL | NULL |
